# Pre- and postsynaptically expressed spiking-timing-dependent plasticity contribute differentially to neuronal learning

**DOI:** 10.1101/450825

**Authors:** Beatriz E. P. Mizusaki, Sally S. Y. Li, Rui Ponte Costa, P. Jesper Sjöström

## Abstract

A plethora of experimental studies have shown that long-term plasticity can be expressed pre- or postsynaptically depending on a range of factors such as developmental stage, synapse type, and activity patterns. The functional consequences of this diversity are unknown. However, in models of neuronal learning, long-term synaptic plasticity is implemented as changes in connective weights. Whereas postsynaptic expression of plasticity predominantly affects synaptic response amplitude, presynaptic expression alters both synaptic response amplitude and short-term dynamics. In other words, the consideration of long-term plasticity as a fixed change in amplitude corresponds more closely to post- than to presynaptic expression, which means theoretical outcomes based on this choice of implementation may have a postsynaptic bias. To explore the functional implications of the diversity of expression of long-term synaptic plasticity, we modelled spike-timing-dependent plasticity (STDP) such that it was expressed either pre- or postsynaptically, or both. We tested pair-based standard STDP models and a biologically tuned triplet STDP model, and investigated the outcome in a feed-forward setting, with two different learning schemes: either inputs were triggered at different latencies, or a subset of inputs were temporally correlated. Across different STDP models and learning paradigms, we found that presynaptic changes adjusted the speed of learning, while postsynaptic expression was better at regulating spike timing and frequency. When combining both expression loci, postsynaptic changes amplified the response range, while presynaptic plasticity maintained control over postsynaptic firing rates, potentially providing a form of activity homeostasis. Our findings highlight how the seemingly innocuous choice of implementing synaptic plasticity by direct weight modification may unwittingly introduce a postsynaptic bias in modelling outcomes. We conclude that pre- and postsynaptically expressed plasticity are not interchangeable, but enable complimentary functions.

**Author summary:** Differences between functional properties of pre- or postsynaptically expressed long-term plasticity have not yet been explored in much detail. In this paper, we used minimalist models of STDP with different expression loci, in search of fundamental functional consequences. Presynaptic expression acts mostly on neurotransmitter release, thereby altering short-term synaptic dynamics, whereas postsynaptic expression affects mainly synaptic gain. We compared cases where plasticity was expressed presynaptically, postsynaptically, or both. We found that postsynaptic plasticity was more effective at changing response times, while both pre- and postsynaptic plasticity were similarly capable of detecting correlated inputs. A model with biologically tuned expression of plasticity also achieved this separation over a range of frequencies without the need of external competitive mechanisms. Postsynaptic spiking frequency was not directly affected by presynaptic plasticity of short-term plasticity alone, however in combination with a postsynaptic component, it helped restrain positive feedback, contributing to activity homeostasis. In conclusion, expression locus may determine distinct coding schemes while also keeping activity within bounds. Our findings highlight the importance of correctly implementing expression of plasticity in modelling, since the locus of expression may affect functional outcomes in simulations.

## Introduction

Learning and memory in the brain, as well as refinement of neuronal circuits and receptive fields during development, are widely attributed to long-term synaptic plasticity [1]. While this notion is not yet formally experimentally proven [2], it has in recent years received strong experimental support in several brain regions, in particular the amygdala [3] and the cerebellum [4]. The notion that synaptic plasticity underlies memory is typically attributed to Hebb [5], but it is in actuality an idea that extends considerably farther back in time, e.g. to Ramon y Cajal and William James [6].

After the discovery by Bliss and Lømo [7] of the electrophysiological counterpart of Hebb’s postulate, now known as long-term potentiation (LTP), much effort has been focused on establishing the induction and expression mechanisms of long-term plasticity. In the 1990s, this lead to a heated debate on the precise locus of expression of LTP, with some arguing for postsynaptic expression, whereas others were in favour of a presynaptic locus of LTP [8]. Beginning in the early 2000’s, this controversy was gradually resolved by the realisation that plasticity depends critically on several factors, notably animal age, induction protocol, and precise brain region [9–11]. Indeed, this resolution has now been developed to the point that it is currently widely accepted that e.g. specific interneuron types have dramatically different forms of long-term plasticity [12, 13], meaning that long-term plasticity in fact depends on the particular synapse type [14]. In retrospect, it is probably not all that surprising that LTP in different circuits is expressed either pre- or postsynaptically, or both, given the diversity of computational functions of different synapses [15]. Nevertheless, the precise functional benefits of having LTP be expressed on one side of the synapse or the other have remained quite poorly explored, with only a handful of classical theoretical papers addressing this point [16–21].

Going back several decades, a multitude of highly influential computer models of neocortical learning and development have been proposed, some of them focusing on aspects such as the rate-dependence of induction [22–24], while others have emphasised the role of the relative millisecond timing of spikes in connected cells [25–27], and some yet have included both [28]. Irrespective of whether timing, rate, or other factors are used to determine the outcome of plasticity in theoretical models, it has virtually always been the case that – with a few notable exceptions [18, 19, 21] – the expression of plasticity itself has been regarded as a simple change in the magnitude of synaptic inputs between neurons of the network. In the absence of better information, this is of course a perfectly reasonable approach, as it is a parsimonious assumption that induction of long-term plasticity manifests itself in the alteration of connectivity weights.

However, the expression of plasticity is not always well modelled by this simple change of instantaneous magnitude. This is because presynaptically expressed plasticity leads to changes in synaptic dynamics, whereas postsynaptic expression does not (Fig.1B). For instance, during high-frequency bursting, the readily releasable pool of vesicles runs out, leading to short-term depression of synaptic efficacy [29], while at some synapse types short-term facilitation dominates [30]. Such short-term plasticity is important from a functional point of view because it leads to filtering of the information that is transmitted by a synapse [31–33]. Short-term depressing connections are most likely to elicit postsynaptic spikes due to brief non-sustained epochs of activity, whereas facilitating synapses require that presynaptic activity be maintained for some period of time before postsynaptic spikes are elicited. In other words, short-term facilitating connections act as high-pass filtering burst detectors [34,35], while short-term depression provides low-pass filtering inputs suitable for correlation detection and automatic gain-control [36–38]. As a corollary, it follows that presynaptic expression of plasticity may change the computational properties of a given synaptic connection. In this case, increasing the probability of release by LTP induction will lead to more prominent short-term depression due to readily-releasable pool depletion, and as a consequence to a gradual bias towards correlation detection at the expense of burst detection [39, 40].

**Fig 1.**
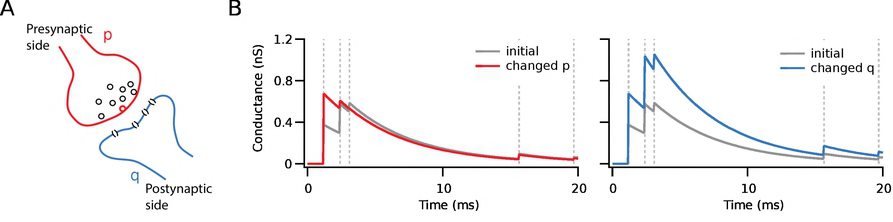
Postsynaptic response to the same stimulus after plasticity depends on expression loci. A - Representation of pre- (red) and postsynaptic (blue) sides of a synapse. B - Initial responses are illustrated in grey, while potentiated ones are in colour. In this example, the amplitude of the first response after learning was set to be the same after both pre- (red) and postsynaptic (blue) potentiation. Whereas with postsynaptic potentiation the gain was increased by the same amount for all responses in a high-frequency burst, with presynaptic potentiation the efficacy of the response train was shifted toward the beginning, enhancing the first response but resulting in no changes over the summed input.

It is long known that the induction of neocortical long-term plasticity may alter short-term depression [16, 41]. While the functional consequences of short-term plasticity itself are quite well described [39, 42], the theoretical implications of *changes* in short-term plasticity due to the induction of long-term plasticity are less well described. Yet, as outlined above, the vast majority of theoretical studies of long-term plasticity assumes that synaptic amplitudes, but not synaptic dynamics, are altered by cellular learning rules. One of the motivations of our present study is the observation that this seemingly innocuous assumption may not be neutral, but in effect a bias, because changing weights in theoretical models of long-term plasticity is equivalent to assuming that synaptic plasticity is solely postsynaptically expressed. This begs the question: What are the functional implications of pre- versus postsynaptically expressed long-term plasticity? Providing answers to this central issue is important for understanding brain functioning, as well as for knowing when weight-only changes in computer modelling is warranted, and when it is not.

Here, we use computational modelling to explore the consequences of expressing plasticity pre- or postsynaptically in a single neuron under two simple paradigms Fig. 2), a time-locked stimulus [26, 43, 44] or the detection of a rate-correlated stimulus [45–47]. Initially, we compare and contrast relatively artificial scenarios, for which the locus of expression is either solely presynaptic, solely postsynaptic, or equally divided between both sides. We then move on to investigating the functional impact of a model with separate pre- and postsynaptic components that were tuned to biological data from connections between neocortical layer-5 pyramidal cells.

**Fig 2.**
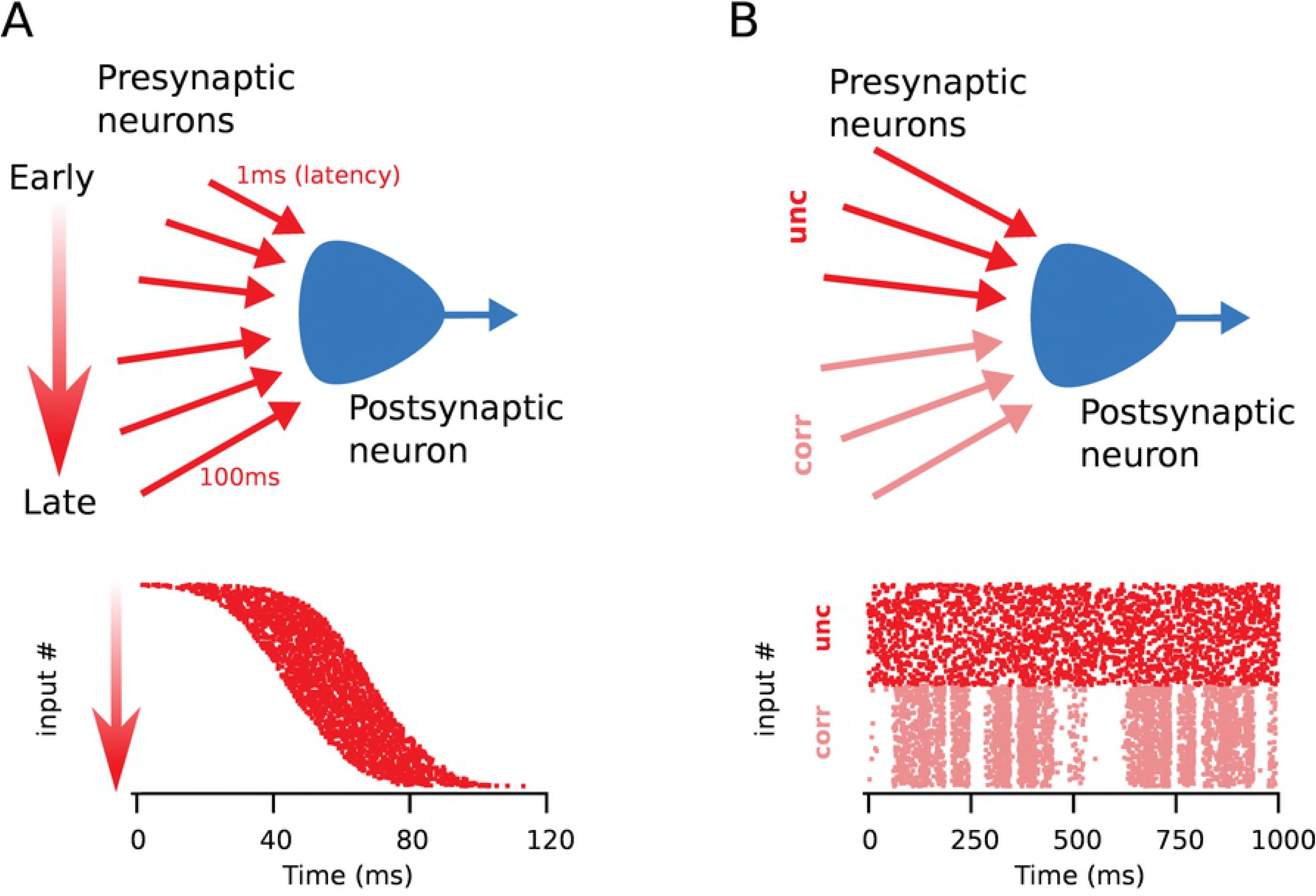
Two different stimulation paradigms were explored. A - Latency reduction: inputs from each presynaptic neuron arrived at different relative times in the volley according to an assigned delay. B - Correlated inputs: half of the inputs had correlated activity, while the rest were independent.

## Results

### Presynaptic expression modelled as changes in stochastic release

For the first set of simulations, we explored the simplest possible differentiation between pre- and postsynaptic efficacy changes. Here, we considered the probability of vesicle release (*P_j_*) and the quantal amplitude (*q_j_*) pre- and postsynaptic quantities respectively. Plasticity was expressed either exclusively on each side or equally divided between both sides.

Since postsynaptic activity depends on the average input across many synapses, one might expect that any differences between pre- or postsynaptic changes should vanish over longer intervals of time. In agreement with this view, there was in this case no appreciable difference between average results. The mean latency shifts (Fig. 3A and B), as well as the decrease of postsynaptic activity duration and increase of postsynaptic firing frequency (respectively Figs. 3D and 3E) did not differ appreciably depending on the implementation. In both cases, sharpening of response only took place after latency reduction was stabilised because frequency and duration were affected by inputs with delays that fell outside of the STDP time window (Fig. 3C). However, in comparison to the purely postsynaptic case, simulations with presynaptic plasticity presented a smaller variance of the latency shift (Fig. 3B), and potentiation developed faster (Fig. 3F). Conversely, depression was slower as the probability of plasticity tended to decrease (Fig. 3F).

**Fig 3.**
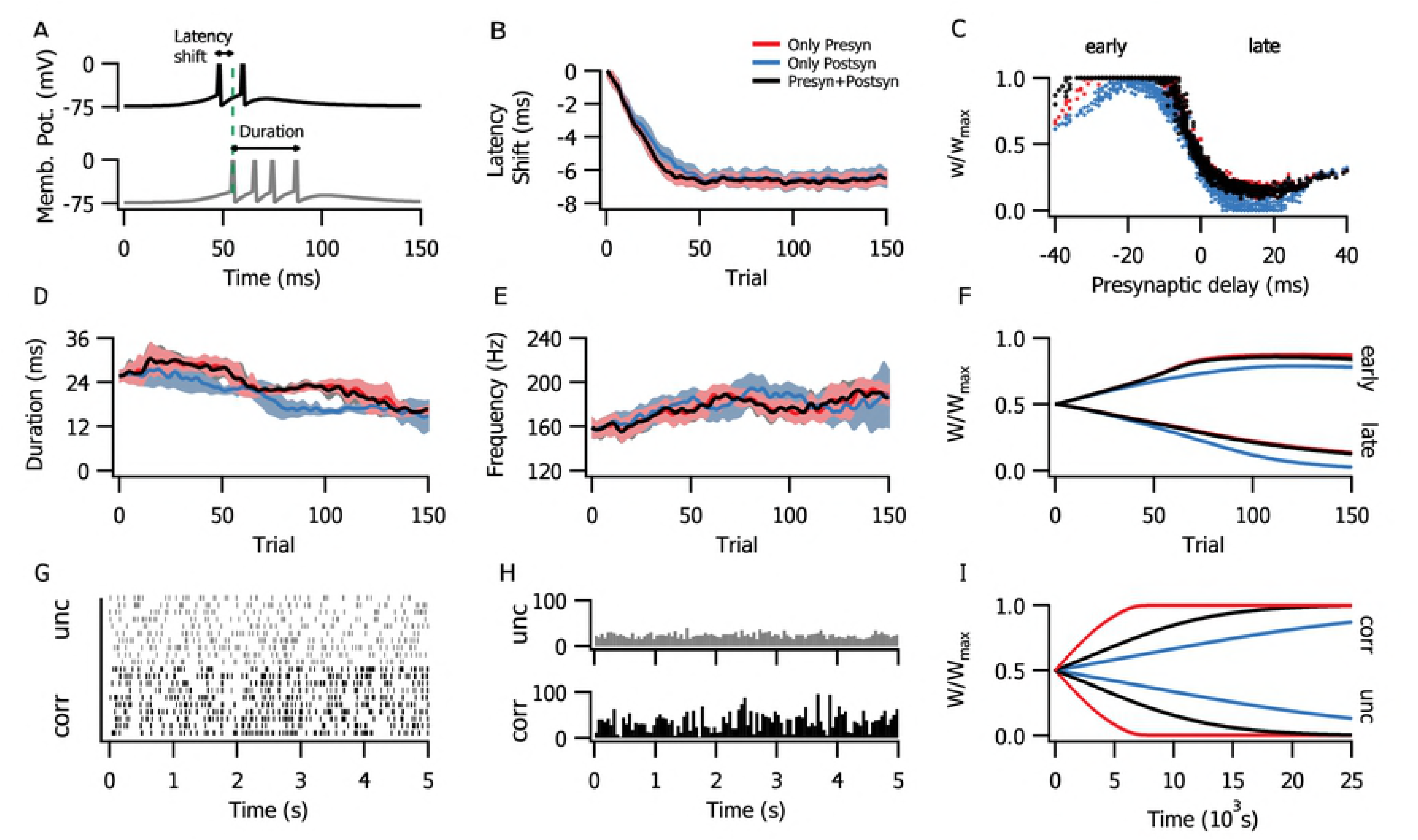
Change of stochastic neurotransmitter release probability lead to faster presynaptic plasticity. The graphs here and below are colour-coded: only presynaptic plasticity (red), only postsynaptic plasticity (blue), or simultaneous preand postsynaptic plasticity (black) are implemented. A-F: Latency reduction configuration. A - Example traces of the postsynaptic membrane potential before (grey) and after (black) plasticity. Initial latency of response is marked by a green dashed line. B - Shortening of postsynaptic latency to spike in comparison to the initial state. C Synaptic weight distribution after 200 trials, normalized for each kind of plasticity expression and sorted by the fixed presynaptic delay. D - Postsynaptic response duration (interval between first and last spike in each trial). E - Postsynaptic intraburst frequency. F - Potentiation of average synaptic weight among early presynaptic inputs (i.e. that arrived within the first half of the stimulus). G-I: Correlated input configuration. G - Input rasterplot sample: correlated inputs shown in black and uncorrelated in gray. H - Histogram of total correlated and uncorrelated presynaptic activities. I - Potentiation of the average synaptic weight among correlated inputs.

In the correlated stimuli paradigm (Fig. 3G and H), potentiation was similarly faster with presynaptic expression of plasticity (Fig. 3I). This happened because plasticity was triggered only after a signal was transmitted, which in the presynaptic case resulted in a positive feedback loop as the probability of potentiation was higher for a more potentiated synapse. This is a consequence of potentiation requiring glutamate release, so that in a high-p synapse, there is an intrinsic propensity for more potentiation. Conversely, depression was slower as the probability of plasticity tended to decrease (Fig. 3F).

### Presynaptic expression modelled as changes in short-term plasticity

We next explored the effects of altering short-term plasticity. This adds another degree of biological realism, since short-term plasticity takes into account the history of presynaptic activity [16]. In this scenario, presynaptic changes generate redistribution of synaptic resources used over a certain time period, instead of an overall amplification. Even if the amplitude of an individual EPSP were affected equally by pre- or by postsynaptically expressed plasticity, the total input from a burst would still differ dramatically depending on the site of expression (Fig. 1B).

Correspondingly, in this case, results differed considerably depending on the specific locus of plasticity in the latency configuration. Postsynaptic expression alone provided the largest latency reduction, and also achieved it faster than the other plasticity implementations (Fig. 4B). Presynaptic expression results were subtle and a defined minimal effect compared to the mixed setting with both pre- and postsynaptic expression. Effects of postsynaptic plasticity over response duration and intraburst frequency (Figs. 4C and 4D) were also more marked. This was a direct result of the increase in total input per burst.

Nevertheless, synaptic efficacy was still potentiated faster and depressed slower in the presynaptic case (Fig. 1E). This was similar to the stochastic case, although less pronounced. This means that even if the rate of learning was effectively faster, presynaptic expression affected timing less efficiently than postsynaptic expression did (Fig. 1F).

**Fig 4.**
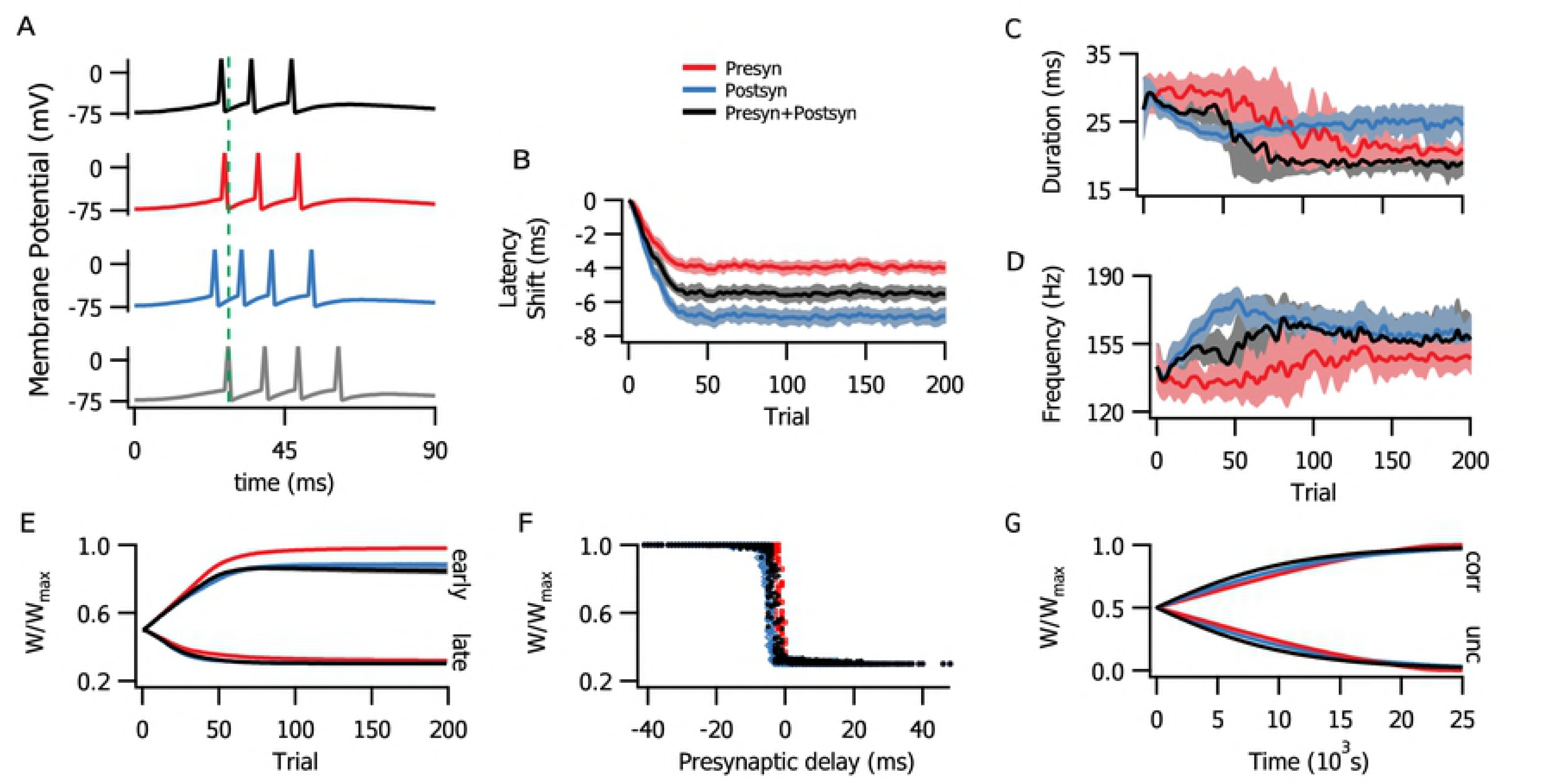
Change of short-term plasticity was less efficient at reducing latency. Graphs here and in subsequent figures are colour-coded: red denotes presynaptic plasticity alone, blue postsynaptic plasticity alone, and black combined pre- and postsynaptic plasticity. A - Example traces of postsynaptic activity before (grey) and after plasticity (coloured). Initial response latency is illustrated by a vertical dashed line. B - Latency reduction was more marked for postsynaptic (blue) than for presynaptic (red) or combined (black) plasticity. C - Combined and presynaptic plasticity reduced response duration better than with postsynaptic expression. D Burst frequency was similarly increased with all three forms of plasticity, although rate change was faster with postsynaptic plasticity. E - Time course of average synaptic weights for early (left) and late (right) inputs. F - Normalized synaptic weight distribution, according to presynaptic delay. G - Time course of average synaptic weights for correlated (left, “corr”) and uncorrelated (right, “unc”) inputs.

On the other hand, plasticity rates in the correlated inputs paradigm evolved differently compared to the stochastic case (Fig. 1G), even though the overall effect on the covariance between pre- and postsynaptic activity was similar (not shown). With depression acting via short-term plasticity, the postsynaptic rate of change was slightly faster than the presynaptic change.

### Contextualization with a biologically tuned model

For an improved biological plausibility, we investigated the interplay between pre- and postsynaptic plasticity in a model that was fitted to data from rodent V1 pyramidal neurons [19, 48]. To isolate the effects of each component, we simply blocked either pre- or postsynaptic changes instead of normalising the total synaptic change in each side, so as to not disrupt of the parameter tuning. We still found that both pre- and postsynaptic plasticity components independently lead to the shortening of postsynaptic latency (Figs. 5B and 5C). As with the above, more abstract modelling scenarios, postsynaptic changes appeared to be more effective at affecting spike timing. When both pre- and postsynaptic plasticity were active, the presence of postsynaptic potentiation enhanced the response compared to presynaptic plasticity alone.

As postsynaptic plasticity in the tuned model lacked the capacity to depress [49], it also lead to inflated postsynaptic frequency and duration if implemented alone (Figs. 5D and E). However, the inclusion of presynaptic LTD was enough to avoid saturation, and the whole model was able to produce a sharpened response. In this case, postsynaptic changes developed faster (Fig. 5F) because of increased postsynaptic frequency.

**Fig 5.**
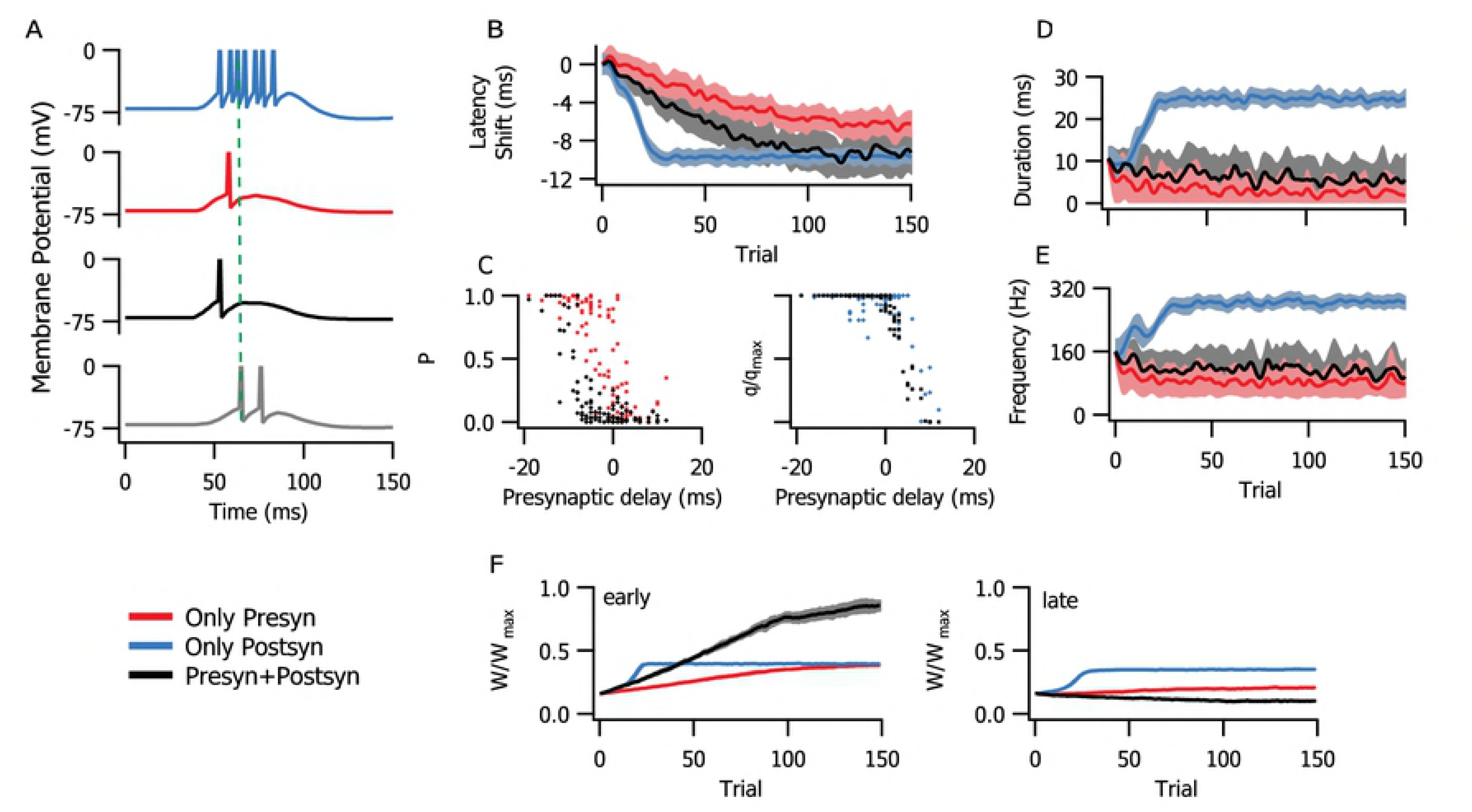
Physiologically tuned model combined pre- and postsynaptic LTP for latency reduction with a controlled activity. The following graphs are colour-coded: only presynaptic plasticity components (red), only postsynaptic component (blue), or both pre- and postsynaptic components (black). A - Example traces of postynaptic activity before (grey) and after plasticity. Initial latency of response is marked by a green dashed line. B - Shortening of postsynaptic latency in comparison to the initial state. C - Distribution of pre- and postsynaptic efficacies after 250 trials. D and E - Duration and intraburst frequency of postsynaptic activity. F Potentiation of average synaptic weight of early (left) and late (right) presynaptic inputs.

In the second configuration, we observed a separation between synaptic efficacy of correlated and uncorrelated inputs (Fig. 6A) without the need of added mechanism of competition [46, 50]. This only occurred when both pre- and postsynaptic components were implemented. This is not achieved through other models with physiologically compatible parameters [47]. We quantified this capacity with a linear separator for the average and variance of *p* values (Fig. 6B).

**Fig 6.**
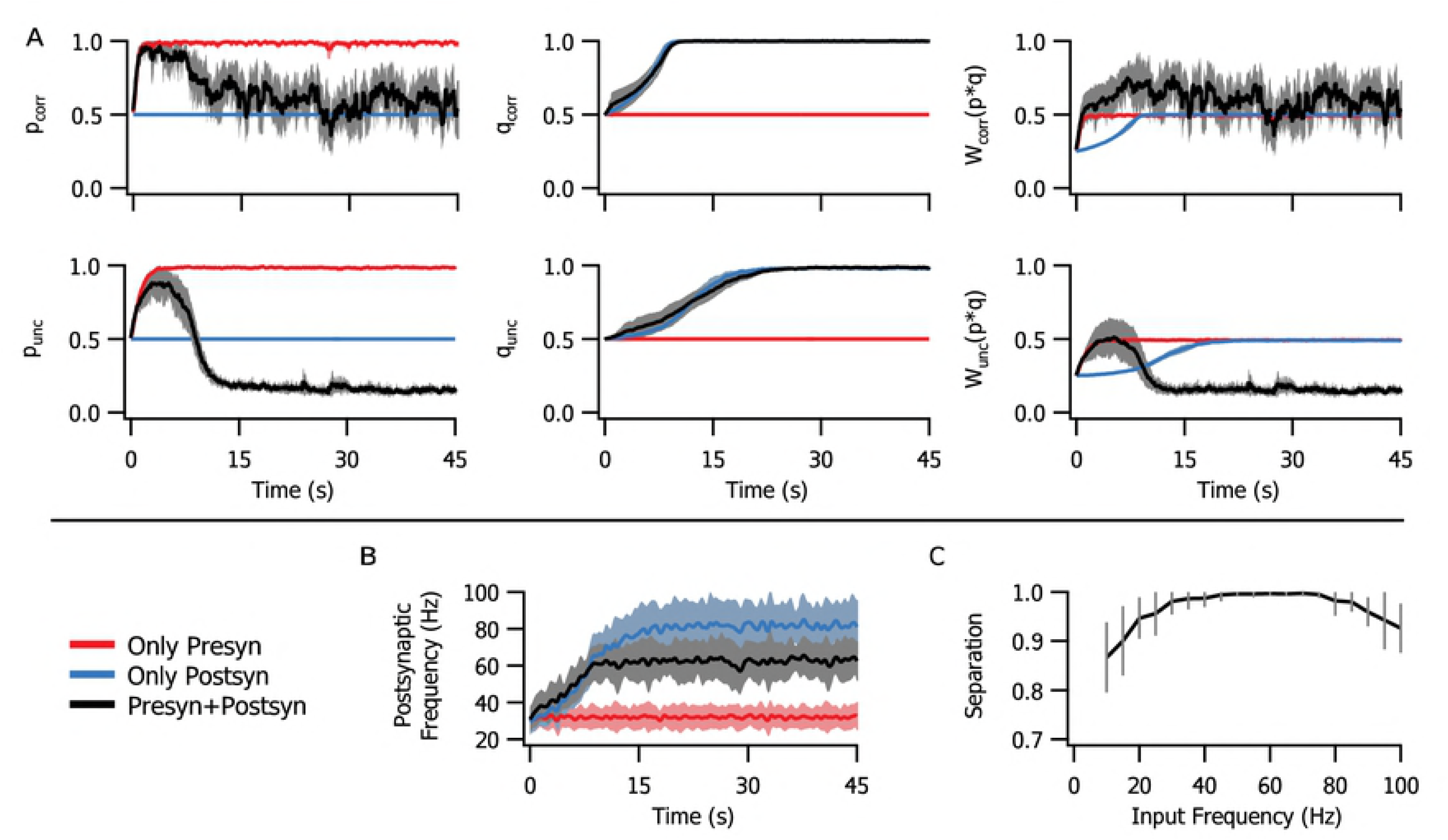
The physiologically tuned model was able to separate between correlated and uncorrelated inputs, but only when there is simultaneous pre- and postsynaptic plasticity. The following graphs are colour-coded: only presynaptic plasticity component (red), only postsynaptic component (blue), or both pre- and postsynaptic components (black). A - Normalized average pre- (p), post- (q) and combined (W) synaptic efficacies of correlated (corr) and uncorrelated (unc) inputs. B - Postsynaptic spiking frequency. C - Average separation between correlated and uncorrelated inputs as a function of presynaptic frequency.

The presynaptic frequency range for optimal separation was between 50 and 80 Hz. On the lower end it was bounded by the correlation time scale, as interspike intervals longer than 20 ms were unable to represent the minimal interval of correlation. On the other end, higher presynaptic frequency yielded overall potentiation that included uncorrelated inputs, limiting the separation from the more potentiated correlated population (see appendix).

In the same way as in the latency configuration, postsynaptic potentiation directly increased postsynaptic firing rate, however in this case the presynaptic component produced no such effect. In combination with postsynaptic plasticity, presynaptic plasticity performed a kind of output control, as its introduction helped to maintain a lower postsynaptic frequency even if *q* saturated (Fig. 6C).

## Discussion

In recent years, it has become eminently clear that diversity in LTP expression is both ubiquitous and considerable, depending on factors such as animal age, induction protocol, and precise brain region [9–11, 15]. In this work, we explored a few possible functional consequences of pre- or postsynaptic locus of plasticity expression, and found that even in a single neuron scenario overall dynamics may be affected by it. Plasticity has in the typical phenomenological model been implemented as a straightforward change in synaptic weight amplitude [26, 51, 52], although there are a few notable exceptions [16–18, 53, 54]. In other words, in the absence of better information, a standard assumption has been that that locus of expression does not matter appreciably for the modelling scenario at hand. Our findings thus challenge this standard assumption, highlighting when it is valid, and when it is not.

We investigated two different learning paradigms, one with differently timed inputs, in which postsynaptic latency to spike was used as a learning measurement, and another under constant stimulation, where a subset of inputs were correlated and potentiated together. We worked with simplified conceptual models, first a simple stochastic STDP implementation and later a more realistic, biologically tuned model of long-term plasticity at in which pre- and postsynaptic components were fitted to connections between neocortical layer 5 pyramidal cells [19].

Our study showed that the locus of expression of plasticity determined affinity for different coding schemes. Presynaptic plasticity as the regulation of flat release probability alone did not result in any differences over average postsynaptic activity measurements compared to the usual postsynaptic expression. However, in the presence of short-term synaptic modulation, presynaptic changes had a milder impact on the timing of postsynaptic spiking output in comparison to changes in quantal amplitude. This was because, as amplitude becomes higher, fewer inputs are needed to evoke a postsynaptic spike, while with presynaptic changes the spike still depends on the sum of a larger number of stimuli. Additionally, overall weight changes developed faster with presynaptic plasticity, in effect upregulating the speed of learning. This effect, however, was not present in the correlated input configuration where both pre- and postsynaptically expressed cases performed similarly.

One could argue that presynaptic short-term plasticity alone was not suitable for postsynaptic rate coding, as it did not affect the average summed input and thus postsynaptic firing frequency remained unchanged. Therefore, in agreement with previously published interpretations [40], presynaptic plasticity appeared to act as a limiter or a form of homeostasis for postsynaptic activity. It is in other words possible that, in some cases, the interpretation of plasticity as a purely postsynaptic phenomenon might lead to overestimation of its effects on postsynaptic firing frequency.

Similar properties were observed in the biologically tuned model with simultaneous pre- and postsynaptic plasticity. Learning results were dramatically affected by postsynaptic plasticity, while the presynaptic side appeared to act more on the rate of learning and on weight dynamics. It is possible that these results could be modified according to the ratio of pre- versus postsynaptic forms of plasticity, thus being optimized according to the computational task at hand. It is noteworthy that the biologically tuned model was also capable of separating groups of correlated and uncorrelated inputs without the need for a competitive mechanism, in an optimal range of input frequencies that depended on input frequency and correlation times.

Since it is possible to specifically block pre- or postsynaptic STDP pharmacologically [41, 49], several of our findings related to the locus of expression of plasticity are experimentally testable. For example, at connections between neocortical layer-5 pyramidal cells, it is possible to block nitric oxide signalling to abolish pre- but not postsynaptic expression of LTP [49]. It is also possible to use GluN2B-specific blockers such as ifenprodil or Ro25-6581 to block presynaptic NMDA receptors necessary for presynaptically expressed LTD without affecting postsynaptic NMDA receptors that are needed for LTP [41, 55]. As a proxy for learning rate, one could explore *in vitro* how blockade of different forms of plasticity expression impacts the number of pairings required for plasticity, or alternatively how the magnitude of plasticity is affected for a given number of pairings [49, 52]. *In vivo*, the impact on cortical receptive fields could similarly be explored. For example, we predict that receptive field discriminability is poorer when presynaptic LTP is abolished by nitric oxide signalling blockade [19].

In conclusion, we challenged the standard assumption of modelling synaptic plasticity as a straightforward weight change by considering plasticity as pre- or postsynaptically expressed, or both. As our collective understanding of LTP expression improves, it is important to understand its overall consequences on circuit dynamics and global functioning of neural networks [56]. We found that even in a simple feed-forward network, the locus of expression could have considerable impact on learning outcome. We speculate that the effect will only be greater in recurrent networks, where presynaptic plasticity at loops and re-entrant pathways will exacerbate the effects of changes in synaptic dynamics due to alterations of the accumulated difference. This additional level of complexity may in particular complicate very large recurrent network models [57, 58]. As the locus of expression of long-term plasticity has been relatively poorly studied, our study highlights the general need for more detailed modelling of the role of the site of expression. In modelling long-term plasticity, correctly implementing changes in weight is thus a matter of gravity.

## Methods

### Neuron model

All of the simulations consisted of one postsynaptic neuron receiving a number of presynaptic Poisson inputs. In the first section, we used a simple leaky integrate-and-fire model defined by

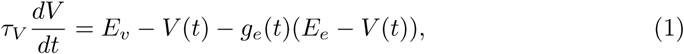

in which the membrane potential *V* decayed exponentially with a time constant of τ_*V*_ = 20ms to the resting value of *E_v_* = −74 mV, and the threshold for an action potential was *V_th_* = −54 mV. After each spike it was reset at *V*_0_ = −60 mV with a refractory period of 1 ms.

Inputs were accounted as conductance-based excitatory contributions with reversal potential *E_e_* = 0 mV, amplitude *q_j_*, summed after the *l^th^* spike of presynaptic neuron *j*, that decayed exponentially with a time constant of τ_*g*_ = 5ms:

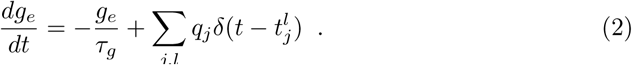

In the the last section, we used the adaptative exponential integrate-and-fire model [59] for increased bursting stability:

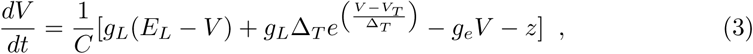

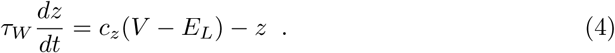

The corresponding parameters for a pyramidal neuron were *C* = 281 pF, *g_L_* = 30 nS, *E_L_* = −70.6 mV, Δ_*T*_ = 2mV, *c* = 4nS, τ_*W*_ = 144ms. Spiking threshold was *V_T_* = −50.4 mV, and after each spike *V* was reset to the resting potential *E_L_* while *z* increased by the quantity *b* = 0.0805 nA (as in [59]).

### Stimulation paradigms

The postsynaptic neuron received either one of two stimulus configurations. The first one was based on [26] and is referred to as latency reduction (Fig. 2A). In every 375-ms-long trial, the postsynaptic cell received a volley of Poisson inputs that arrived with fixed delays, normally distributed around a time reference, per specific presynaptic neuron. Each input lasted for 25 ms with a spiking frequency of 100 Hz. We measured the time to spike of the first postsynaptic spike in response to a bout of stimuli using the mean of the presynaptic delay distribution as a reference point. For clarity, in the Results, curves that represent latency shift, intraburst frequency or burst duration were smoothed using a moving average filter with a window of three points.

The second type of stimulation paradigm was based on [45] (Fig. 2B). This configuration consisted of continuous Poisson inputs with fixed frequency. However, half of the inputs had correlated fluctuations of activity, with a time window of τ_*corr*_ = 20 ms, while the other half was uncorrelated.

### Additive STDP model

For the majority of the simulations we opted to implement STDP with the simple additive model proposed by Song and Abbott [26]:

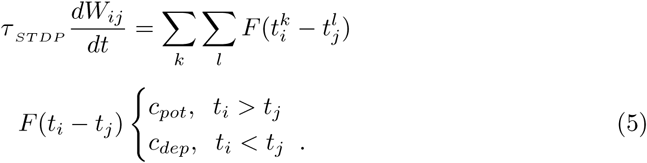

Each increment to the synaptic weights *W_ij_* (since there was only one postsynaptic cell we consider *W_j_* = *W_ij_* for the rest of this paper) was computed after a pair of pre- and postsynaptic spikes, and the parameters were set to τ_*STDP*_ = 20ms, *c_pot_* = 0.005, and *c_dep_* = −0.00525. We separated the synaptic weight *W_j_* as a product between pre- and postsynaptic counterparts, probability of release *P_j_* (0, 1] and quantal amplitude *q_j_* (0, *q_max_*] respectively, so that *W_j_* = *q_j_P_j_*. The probability of release was simulated in two different ways, one by regulating the probability of stochastic interactions and the other by short-term plasticity.

When the weight convergence rates were compared, we had to ensure that Δ*W* =*W^f^* − *W^i^* per time step was the same for all simulations. Therefore, we normalized the chages so that if only *q* was changed:

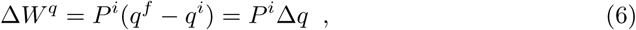

and if only *P* was changed,

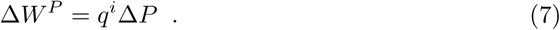

The initial value of all simulations was the same for *P* and *q*, so in these cases Δ*P* = Δ*q* ≡ Δ. This amount was equally divided between *P* and *q* when both were changed simultaneously:

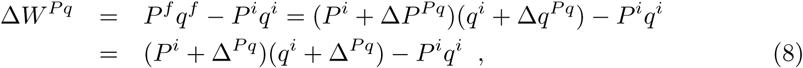

so that

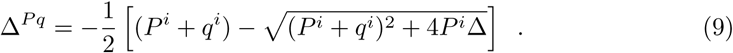

The largest possible change for *P* or *q* separately was Δ_*tot*_ = 1 − *q^i^*. To keep the same range of *W* for changing *P* and *q* simultaneously, we limited the maximal values *P* and *q* in this case at 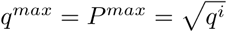.

### Biologically tuned STDP model

We compared the results of the straightforward additive model to a slightly more complex STDP model that acts separately over pre- and postsynaptic factors [19]. Parameters were fitted to experimental data from connections between pyramidal cells from layer 5 of V1 [41, 48, 49]. The equations for pre- and postsynaptic changes followed:

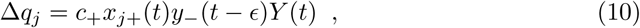

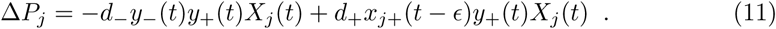

where 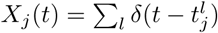 is increased at each spike from the presynaptic neuron *j* and 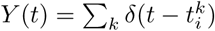 at each spike from the postsynaptic neuron. ∊ is to emphasize that Δ*W* was calculated before *x*_*j*+_ and *y*_−_ were updated, upon the arrival of a new spike. *y*_+_ and *y*_−_ are postsynaptic traces,

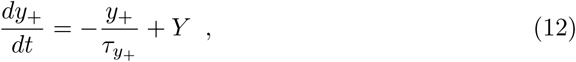

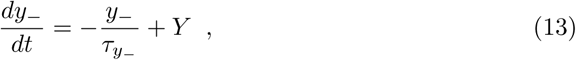

with decay times τ_*y*+_ and τ_*y*−_ respectively, and *x*_*j*+_ was a presynaptic trace with decay time τ_*x*+_:

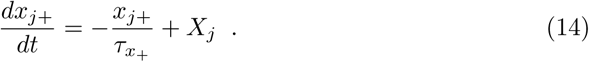

The parameter values were taken from [19]: *d*_−_ = 0.1771, τ_*y*−_ = 32.7ms, *d*_+_ = 0.15480, *c*_+_ = 0.0618, τ_*y*+_ = 230.2ms and τ_*x*+_ = 66.6ms. To avoid manipulation of the fitting, weight changes were not normalized in this case.

In the last section, we used a linear least squares separator to classify presynaptic inputs according to synaptic weight average and variance.

### Presynaptic factor

Presynaptic control of the probability of release per stimulus was implemented either as a Markovian process or as short-term plasticity. In the former case, probability (*P_j_*) of stochastic neurotransmitter vesicle release followed a binomial distribution. Each presynaptic neuron had *N* = 5 release sites that functioned independently. In the second we considered a dynamic modulation of the EPSPs through STP. The probability *P_j_* was decomposed into the product of instantaneous probability of release *p_j_*(*t*) and availability of local resources *r_j_*(*t*), resulting in the following synaptic efficacy:

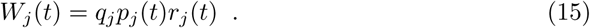

In the latter case, the dynamics of *p_j_*(*t*) and *r_j_*(*t*) followed the model proposed by Tsodyks and Markram [60]:

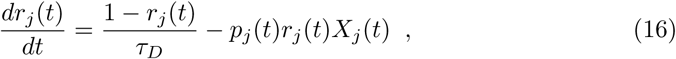

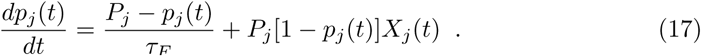

Depression and facilitation time constants, τ_*D*_ = 200 ms and τ_*F*_ = 50 ms respectively, were chosen as representative values for connections between pyramidal neurons [61]. The resulting short-term plasticity could be either depressing, if *P_j_* > *P_C_*, or facilitating, if *P_j_* < *P_C_*. For the values of τ_*D*_ and τ_*F*_ used, *P_C_* ≈ 0.3.

## Supporting information

### S1 Appendix

**Rate model** A simple firing rate model with linear response was able to illustrate how correlated and uncorrelated input separation was achieved without competition between the two populations. Considering independent Poisson inputs with fixed firing rate we found there is a determined, non-zero average value for *P*. The LTP model (eqs. 10 and 11) was converted to time-averaged values:

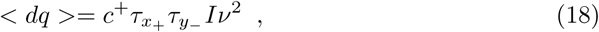

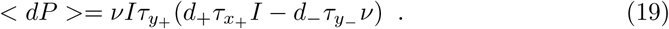

Postsynaptic output ν was then considered as a simple firing rate model with linear relation to average input *I*, weighted by average synaptic efficacy (eq. 15):

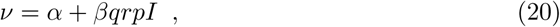

where α and β (for fixed values of *q* and *P*) were fitted to data from simulations without plasticity. Since *I* was fixed, we could consider stationary values for *r*(*t*) and *p*(*t*), 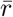 and 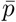, from eqs. 16 and 17:

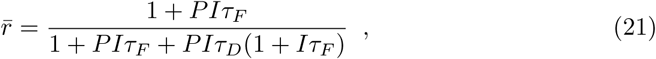

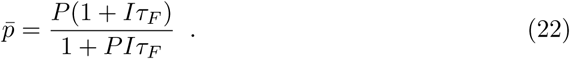

We thus have < *dq* > (*P, q, I*) and < *dP* > (*P, q, I*) for LTP:

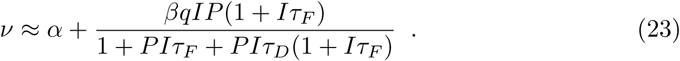

In the vector field *dP* × *dq* (Fig. 7A), it is visible that *P* tends to the a specific value (*P*\*) at the intersection of the higher\**p*-nullcline and the maximum value *q_max_* which corresponds to the average value of *P* for uncorrelated inputs. This is in contrast to correlated inputs, which tend to potentiate to the maximum. This value tends to increase with frequency, limiting the range of separation of the more potentiated correlated population.

**Fig 7.**
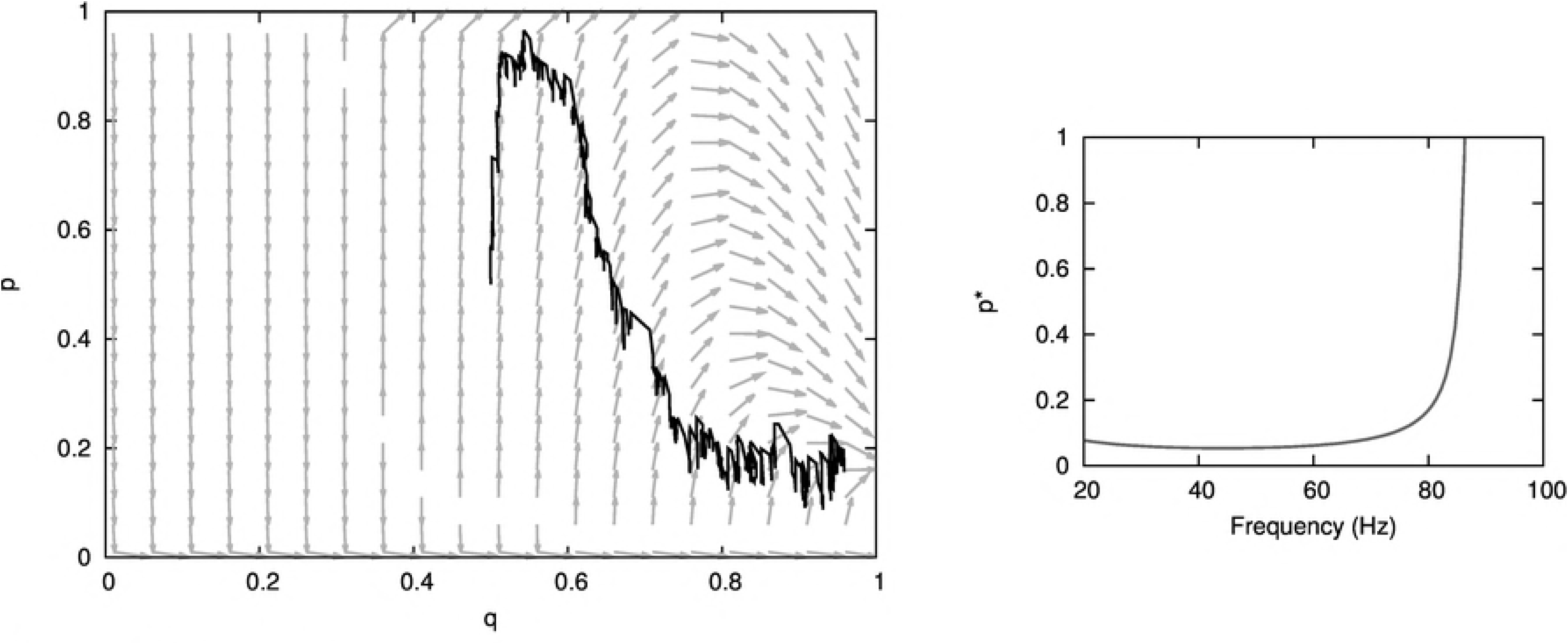
**S1** A - Rate model *p* × *q* vector field for uncorrelated synaptic inputs. The lines show corresponding simulation averages for uncorrelated (yellow) and correlated (gray) inputs. B - *Point of convergence *P*\* as a function of presynaptic frequency.

## Data availability statement

All files will be available on GitHub.

## Acknowledgements

We thank Alanna Watt and Mark van Rossum for suggestions, help, and useful discussions.

## Author contributions

**Conceptualization:** Beatriz E.P. Mizusaki, Rui P. Costa, P. Jesper Sjöström.

**Formal analysis:** Beatriz E.P. Mizusaki, Sally S.Y. Li.

**Funding acquisition:** Beatriz E.P. Mizusaki, Sally S.Y. Li, Rui P. Costa, P. Jesper Sjöström.

**Investigation:** Beatriz E.P. Mizusaki, Sally S.Y. Li.

**Methodology:** Beatriz E.P. Mizusaki, Rui P. Costa.

**Project administration:** P. Jesper Sjöström.

**Resources:** Rui P. Costa, P. Jesper Sjöström.

**Software:** Beatriz E.P. Mizusaki, Sally S.Y. Li, Rui P. Costa.

**Supervision:** P. Jesper Sjöström.

**Validation:** Beatriz E.P. Mizusaki.

**Visualization:** Beatriz E.P. Mizusaki.

**Writing – original draft:** Beatriz E.P. Mizusaki, Rui P. Costa, P. Jesper Sjöström.

**Writing – review & editing:** Beatriz E.P. Mizusaki, Rui P. Costa, P. Jesper Sjöström.

